# Turing-like patterns can arise from purely bioelectric mechanisms

**DOI:** 10.1101/336461

**Authors:** Micah Brodsky

## Abstract

Turing-like patterns can potentially occur in non-neural (non-excitable) tissues through strictly bioelectric processes, without involving transcriptional (gene) regulation, cell migration, or traditional reaction-diffusion mechanisms. Small molecules that gate transmembrane ion channels are often charged and capable of passing through intercellular gap junctions, and their transport under the influence of trans-junction electric fields furnishes a bioelectric feedback loop. We develop an analytically tractable, circuit-based model of this phenomenon and show that it can lead to spontaneous formation of spatial patterns in ligand density and membrane voltage under physiologically plausible conditions. The process is distinct from Turing’s reaction-diffusion paradigm but closely analogous to the spontaneous formation of patterns in colonies of chemotactic bacteria.

## Introduction

The divergent mechanisms employed in development to pattern undifferentiated tissues can appear bewildering in variety. Two principal classes stand out, however, which might be termed “feed-forward patterning” and “feed-back patterning”. Feed-forward processes, as anticipated in Wolpert’s classic theory of positional information and exemplified by the drosophila segmentation cascade, employ positional information encoded in maternally-provided pre-patterns, boundary conditions, or planar polarizations, and compute a cascade of spatial functions based on this input. In the steady state limit, none of these functions require any memory, and as there are no cross-stage cycles in the computation, their combination can be memoryless as well. Perturbations naturally decay with time, and there is no way to generate or reinforce the underlying spatial cues; they must be provided from the outside. Such processes are fast, determinate, and often scale-invariant.

Feed-back processes, on the other hand, such as Turing patterns (Watanabe and Kondo 2015, Marcon and Sharpe 2012, Turing 1952), are capable of generating their own positional information, either through instabilities or by bootstrapping it from weakly asymmetric initial conditions. In their simplest form, they employ tight spatial feedbacks, positive at short ranges but negative at longer ranges – the famed “local activation, lateral inhibition” criteria – so as to select one or more spectral wavelengths for spontaneous amplification (Rabinovich, Ezersky, and Weidman 2000). As a result, they have baked-in length scales associated with their instabilities and are often indeterminate, changing behavior on differing sizes and shapes of substrate. Owing to their mathematical similarities, widely divergent feedback processes can produce remarkably similar-looking results (Hiscock and Megason 2015).

In general, feedback processes are not particularly fast and may fall victim to metastability. They are, however, highly capable in self-repair and might serve as a foundation for tissue homeostasis. In earlier theoretical work (Brodsky 2016), we argued that the simultaneous combination of feed-forward and feed-back patterning has compelling evolutionary advantages, combining the virtues of both, and may serve as a useful paradigm for interpreting the apparent redundancies seen in developmental patterning programs.

Among feed-back processes, the largest share of research attention has gone to the Turing mechanism – reaction-diffusion where the reaction’s stability in a well-mixed environment is broken by differing diffusion rates of the reagents. In proposed biological realizations, however, these “reagents” are often average concentrations of various cell types, and “diffusion” may be the quasi-random motion of cells or even more exotic processes for interaction at a distance such as filopodial contacts (Watanabe and Kondo 2015). Here, we demonstrate the possibility of a rather different feed-back patterning mechanism, a novel bio-electrochemical process based on cell membrane potential and the diffusion of small molecules through gap junctions. Mathematically, it is most closely related to the pattern-forming processes seen in chemotactic aggregation, as in e.g. bacterial colonies (Hillen and Painter 2008, Budrene and Berg 1995). The result can be both electrochemically demarcated “aggregates” as well as complex feed-back patterns reminiscent of Turing patterns.

Gap junctions are pervasive and important in animal development, although their precise roles often remain unclear. In addition to electric currents, they can transmit small molecules such as second messengers, ATP, Ca++, serotonin – species that are very often charged. Thus, the voltage seen across a gap junction – the difference in membrane potential between the apposed cells – can have a major impact on transport rates. Such voltage-mediated transport is thought to play an important role in processes such as left-right axis patterning (Esser et al. 2006). What if, however, the transported chemical species directly regulates membrane voltage, as by gating a ligand-gated ion channel, or perhaps by modulating an ion channel’s expression, e.g. siRNA (Valiunas et al. 2005)? This introduces a new feedback that has been shown to be capable of causing interesting phenomena such as spontaneous polarization (Pietak and Levin 2016). In particular, if the sign of the feedback is positive – a positively charged ligand causing hyperpolarization and thereby increased attraction, or similarly a negatively charged ligand causing depolarization – then the ligand may be capable of spontaneous aggregation, leading to symmetry breaking and pattern formation. The attraction of nearby ligand, and its consequent depletion at longer range, constitutes the critical “local activation, lateral inhibition”. We refer to this phenomenon here as *auto-electrophoresis*.

Using the model developed below, we derive the conditions under which auto-electrophoretic pattern formation will occur. The phenomenology is somewhat reminiscent of the first order phase transition seen when a vapor spontaneously condenses into droplets, including a definite “dew point” threshold that must be crossed for spontaneous nucleation, as well as a more lenient “coexistence region” where existing aggregates will persist but not nucleate on their own.

Not all channel / ligand pairs will be practically capable of auto-electrophoretic pattern formation. One example likely capable of this process are the CNG channels their ligands the cyclic nucleotides cAMP and cGMP. cAMP and cGMP are well-known to be gap junction permeable and are negatively charged at physiologic pH, and several members of the CNG family are capable of causing depolarization with strong, cooperative sensitivity to the ligand (Kaupp and Seifert 2002, Ruiz et al. 1999).

In order to explore these possibilities, one approach would be to conduct exhaustive simulations with a detailed simulator such as BETSE (Pietak and Levin 2016). In a companion work, we follow this strategy, using a genetic algorithm to direct a search through the parameter space, and indeed we demonstrate spot and stripe patterns spontaneously generated using CNG channel models (Brodsky and Levin 2018). Here, however, we show that one can also construct a simplified physical model with far fewer parameters, from which wide-ranging analytical predictions can be directly derived. This model bears strong similarity to the classic Keller-Segel model of bacterial chemotaxis (Hillen and Painter 2008), with membrane voltage taking the role of chemoattractant concentration and intracellular ligand concentration taking the role of bacterial population density. The model is distinct from the classic Turing reaction-diffusion paradigm, instead being an *advection*-diffusion system.

We first derive the model and demonstrate its close parallels with chemotaxis. Subsequently, we derive several formulae directly from the model that predict the circumstances in which spontaneous pattern formation will occur as well as the timescale required and the characteristic size of pattern features, as a function of experimentally measurable quantities. These predictions will be valuable in guiding efforts toward experimental demonstration as well as for identifying possible natural, in vivo instances. We then simulate the model numerically, efficiently computing large simulations that showcase the qualitative characteristics of the auto-electrophoretic patterning process.

## Mathematical model

To begin with, we need a simplified model of a cell. Electrically, we can model each cell as a simple circuit fragment, along the lines of the classic Hodgkin-Huxley models. We assume approximately fixed extracellular and intracellular ion concentrations, represented as constant reversal potentials, and allow these to affect V_mem_ through variable resistances representing channel permeability. Thus, each cell is modeled as follows:

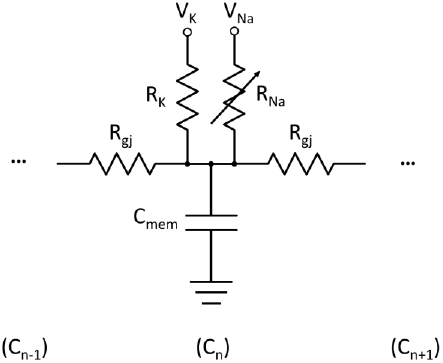

Where cells are arranged one after another in a line representing a one-dimensional tissue. We ignore the effects of lateral cell-to-cell capacitance, which should be minimal for slow phenomena like auto-electrophoresis. One or both of the two transmembrane resistances, R_K_ and R_Na_, is modeled a function of ligand concentration, representing ligand gating. We ignore voltage gating and voltage-dependent GHK transport for now, as their effects on the slow, small-amplitude variations present during early pattern formation can be approximated through modifications to the ion potentials and transmembrane resistances (adjusting them to produce a linearized approximation to the nonlinear channel’s local behavior, which is perfectly adequate for the linear stability analysis employed below). This diagram also illustrates only Na and K affecting membrane potential, but it can capture other ungated ions equally well, lumped together as a Thevenin equivalent resistance. Applied to this system, Kirchoff’s current law gives

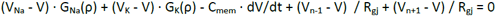

Where G_Na_ = 1 / R_Na_ is the sodium conductance and G_K_ = 1 / R_K_ is the potassium conductance. Without loss of generality we can assume it is G_Na_ that is gated, leaving G_K_ a constant. (The results remain the same if reversed, except the sign the voltage is inverted.) If we restrict our attention to patterns larger than a single cell, we can simplify things further by taking the continuum limit, yielding a partial differential equation:

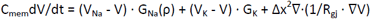

Where Δx is the cell length. This is a form of the diffusion-decay equation, and it generalizes immediately 2d and 3d tissues.

Chemically, the transport of ligand among cells can be modeled by the combination of Fick’s law of diffusion and the Einstein relation for drift under an electric field, a combination known as the Nernst-Planck equation. Discretely, we have

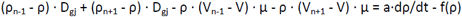

Where ρ is the ligand concentration, D_gj_ is the diffusive conductance between neighboring cells, a is the cell volume, μ = D_gj_q/kT ≈ D_gj_z/26mV is the ligand mobility, z the ligand charge, and f(ρ) models any intracellular production and decay kinetics for the ligand. To be more accurate, one should treat discrete cells as connected via the nonlinear GHK flux equation, which takes into account the effect on the electrophoretic flux of sharp variation in ligand concentration between neighboring cells. However, sharp variations are absent in the continuum limit, and absent even in the discrete case in the early stages of pattern formation as well as for well-developed but long-wavelength patterns, so we can reasonably omit this complication. Thus, in the continuum limit, we have

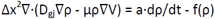

which is the Nernst-Planck equation. We can then transform our continuum limit system of equations into a simpler, non-dimensional form via several substitutions (where italics are used to denote non-dimensionalized quantities). First, we set

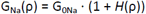

Where G_0Na_ represents the baseline, ungated sodium conductance and *H*(ρ) captures the channel gating behavior, e.g. through a Hill function, representing how many fold the depolarizing conductivity increases above the baseline leakage. For a simple linear model, *H*(ρ) = ρ, and ρ is scaled in units of the concentration that doubles sodium conductance. For more general models, we may find it convenient to scale ρ such that *H*(ρ^0^) = ρ^0^ for some particular steady-state ρ^0^.

We then use the following scaling substitutions:

| <u>Dimensionless scale substitution</u> | <u>Associated scale parameters</u> | <u>Parameter formulae</u> |
| --- | --- | --- |
| $V = (V - V^0) / V^*$ | Voltage scale<br>Voltage midpoint | $V^* = (V_{Na} - V_K) / 2$<br>$V_0 = (V_{Na} + V_K) / 2$ |
| $t = t / \tau_E$ | Electrical timescale<br>Diffusive timescale | $\tau_E = C_{mem} / G_{0Na}$<br>$\tau_D = a / D_{gj}$ |
| $x = x / x_0$ | Length scale | $x_0 = \Delta x / \sqrt{(R_{gj} G_{0Na})}$ |
|  | <u>Dimensionless parameters</u> | <u>Parameter formulae</u> |
| | Leak current ratio | $g = G_K / G_{0Na}$ |
| | Diffusion ratio | $D = \tau_E / \tau_D \cdot (\Delta x / x_0)^4$<br>$= R_{gj}^2 G_{0Na} C_{mem} D_{gj} / a$ |
| | Charge ratio | $Z = -zV^* / V_{th}$<br>($V_{th} = kT / q \approx 26mV$) |
| | Kinetics function | $f = \tau_E / a \cdot f$ |

The electrical timescale τ_E_ represents the characteristic RC relaxation time of the membrane potential through (somewhat arbitrarily) the baseline Na conductance. The diffusive timescale τ_D_ represents the characteristic diffusive relaxation time between neighboring, GJ-connected cells. In typical tissues, the electrical timescale τ_E_ may be on the order of tens to hundreds of ms, while the diffusive timescale τ_D_ may be on the order of minutes to hours. Later, we will also define the slower pattern-formation timescale,

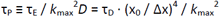

where k_max_ is the leading wavenumber of the pattern.

The key dimensionless parameters above are the leak current ratio, *g*, which determines how much influence gated Na conductance can have on membrane potential, and *Z*, which indicates the strength of the ligand electrostatic interaction. These two parameters, along with ligand concentration and the sharpness of the gating function *H*, determine whether the system will be able to experience the auto-electrophoretic instability.

With these substitutions, the following system of equations results, representing our completed model of auto-electrophoresis:

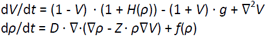

These equations are strikingly similar to the Keller-Segel equations used to model chemotaxis, differing only in the unusual form of the “chemoattractant” V’s production and decay terms, reflecting the fact that V is bounded between V_K_ and V_Na_. The other notable difference is that f(ρ) represents chemical kinetics rather than population dynamics, and therefore can include a zeroth order production term.

### A graphical thermodynamic interpretation

We can gain some basic, physically-motivated insight into phenomenology of auto-electrophoresis with a simple graphical analysis of the above equations, demonstrating a close analogy with the thermodynamics of the vapor/liquid phase transition. The Nernst-Planck equation can be rewritten as the steepest descent of free energy per ligand molecule – i.e., a chemical potential (Zheng and Wei 2011). If we posit an effective chemical potential of the form

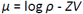

where the first term represents the entropy of dispersal and the second term represents electrostatic energy (with the Z parameter relating the two, playing a role much like the reciprocal of temperature), then our Nernst-Planck equation above can be written

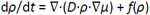

The electrostatic energy is, of course, not the fundamental electrostatic interaction between charged ligand particles, but rather, an effective, attractive interaction due to their influence on membrane potential. From this perspective, ligand auto-electrophoresis is the competition between a short-range attractive force and entropic dispersal. When the “temperature” 1/*Z* is low enough, we may expect to see spontaneous condensation. If we omit chemical kinetics *f*(*ρ*), the analogy can be illustrated by plotting “isotherms” of the chemical potential – curves of *μ* vs. specific volume (1/*ρ*) at fixed *Z* for uniform, steady state solutions:

**Figure 1.**
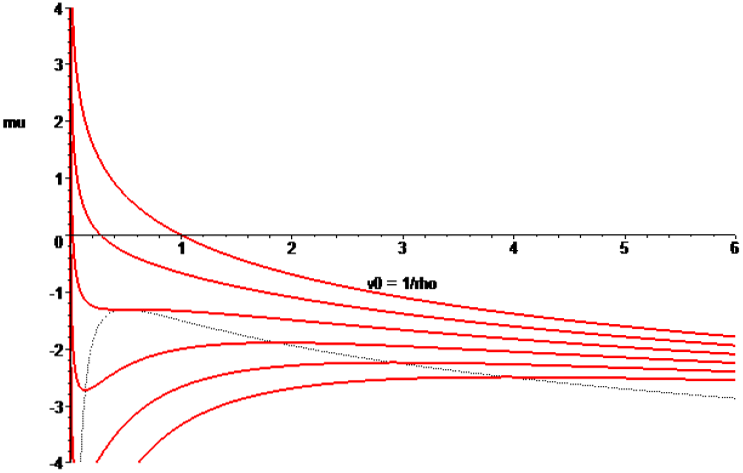
“Isotherms” showing chemical potential (effective free energy per molecule) vs. volume (1 / *ρ*) at fixed Z values. Instability occurs in regions of positive slope (dotted boundary). g = 1, Z = [0, 2, 4, 6, 8, 10] (top to bottom), linear gating.

For sufficiently large *Z*, there appears a region of positive slope (dotted region) – where free energy per particle *decreases* with decreasing volume. Just as in the classical Van der Waals model of the vapor/liquid transition, this represents a regime of thermodynamic instability – where the ligand will spontaneously collapse and separate into two distinct phases, one of high density, one of low density. This region in fact corresponds precisely to the conditions for spontaneous pattern formation derived in subsequent sections. In the simplified case here, with linear channel gating and no chemical kinetics, one can show that the region of instability is

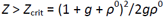

which is plotted as the dotted line above. The smallest possible value of *Z*_crit_ in this case is 2, approached at large *g* and *ρ*^0^. A more general formula is derived in a later section. Immediately surrounding this instability region, one can also predict the existence of a “coexistence region” using the Maxwell equal-area construction, where the two distinct phases will persist if present but will not spontaneously nucleate from a homogeneous mixture – i.e. where a long-wavelength pattern will persist or perhaps nucleate from a finite-amplitude disturbance but will not spontaneously form.

Within the region of thermodynamic instability, a homogeneous, well-mixed solution as plotted above is not actually stable. Instead, we can expect spontaneous condensation of “droplets”, existing in a rough chemical equilibrium with the surrounding “vapor”. The overall ligand density, and hence the relative proportions of the two phases, determines the character of the resulting pattern. Mostly vapor produces a droplet-like spot pattern, while mostly liquid produces a pattern of bubble-like holes, and intermediate concentrations can produce stripe-like, labyrinthine configurations. The characteristic size of these spontaneously condensing pattern features can be determined from linear stability theory (see below).

However, such a pattern of droplets is not necessarily stable either, as the free energy deficit due to the droplet boundaries – “surface tension” – means that smaller droplets are less energetically favorable than larger droplets. Thus, droplets will contract into round shapes, and large droplets will grow at the expense of smaller ones, or droplets may merge together, yielding a coarser and coarser pattern as time goes on. This is a known phenomenon not only in vapor/liquid systems but also in chemotaxis models, where it can be shown to progress on a slow, logarithmic timescale (Painter and Hillen 2011, Hillen and Painter 2008). Such “coarsening” behavior is demonstrated here in the first of the numerical examples below.

Thus far we have ignored chemical kinetics – a non-equilibrium contribution, in the thermodynamic analogy. If ligand is continually produced and destroyed across space, then it is possible converge to a dynamic equilibrium with a stable, fixed-scale pattern. It is also possible to converge to a chaotic attractor with ever-changing patterns – seen under some configurations with nonlinear kinetics. In either case, the character of the pattern – spot, stripe, or net – is ultimately controlled by the relative balance of ligand production and decay, which determines the relative fractions of space occupied by liquid and vapor.

### Quantitative analysis of the model

We now more carefully derive a series of quantitative predictions. The model system has the uniform steady state solution(s):

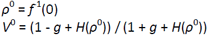

Using standard techniques of linear stability analysis, as in Turing’s original work, one can derive the behavior of small perturbations about the steady state, indicating when then system is capable of spontaneous symmetry breaking and pattern formation. For solutions of the form,

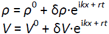

We have

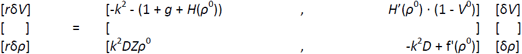

Since *H*(*ρ*) ≥ 0, and for stable chemical steady states, *f*’(*ρ*^0^) ≤ 0, the trace of the matrix is negative, i.e. one or both *r* eigenvalues are negative. Thus, det < 0 indicates pattern-forming instability at wavenumber *k*. I.e.,

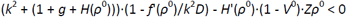

In the absence of chemical kinetics, i.e. *f*’(*ρ*^0^) = 0, this simplifies to

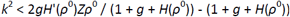

This leads to a simple test for pattern-forming instability:

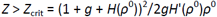

For *Z* values above the critical value, there exists at least one wavelength that is unstable. For a singly charged ligand, typical *Z* values will be in the range of 1.5 to 3 (at the lower end for a nonselective gated channel), and for a doubly charged ligand, 3 to 6. Interestingly, *Z*_crit_ depends only on *g, ρ*^0^, and *H*; the particulars of diffusion, capacitance, and gap junction interconnectivity are irrelevant. They affect the time and length scales of the pattern, but not its presence or absence. For a simple linear gating model (*H*(*ρ*) = *ρ*), the smallest possible value of *Z*_crit_ is 2, achieved at large *g* and *ρ*^0^ – i.e. when ungated conductance of the gated ion (sodium, we’ve assumed) is comparatively small. The plot below illustrates *Z*_crit_ as a function of *g* and *ρ*^0^. For channels with cooperative binding, large Hill exponents n can drive *Z*_crit_ even smaller, by a factor approaching n when far from saturation. (This can be shown by scaling ρ such that *H*(*ρ*^0^) = *ρ*^0^, in which case 0 < *H*’(*ρ*^0^) < n.) Nonzero chemical kinetics inevitably weakens the instability, although it can foster interesting long-term patterning behavior when moderate enough so as not to destroy the instability entirely.

**Figure 2.**
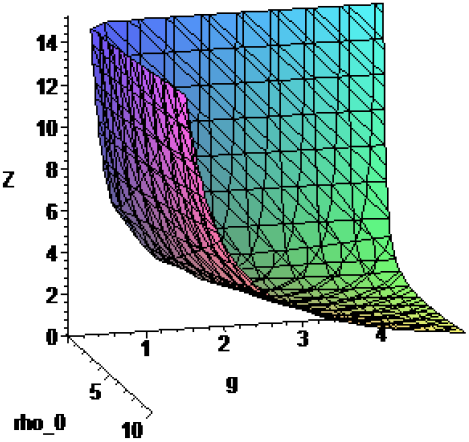
*Z*_crit_ as a function of *g* and *ρ*_*0*_ for linear channel gating.

Noting that the length and time scales have no impact whatsoever on the presence of spontaneous patterning (aside from setting the relevant scale for the effect of chemical kinetics via the *D* parameter), we can make a further approximation. In typical tissues, the electrical timescale τ_E_ is many orders of magnitude faster than the diffusive timescale τ_D_. This large timescale separation means we can approximate the electrical phenomena as instantaneously reaching steady state. The system equations then read,

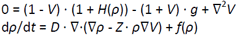

and the linearized equations for small perturbations,

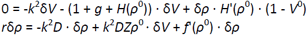

These can be solved to give an expression for the rate of exponential growth as a function of wavelength 2π/*k*:

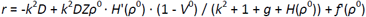

We can use this to derive for any given set of parameters the most unstable wavelength, which will tend to dominate the pattern (assuming it is not already smaller than the physical size of a cell):

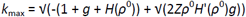

The example of a simple linear gating model with *Z* = 4 is plotted below:

**Figure 3.**
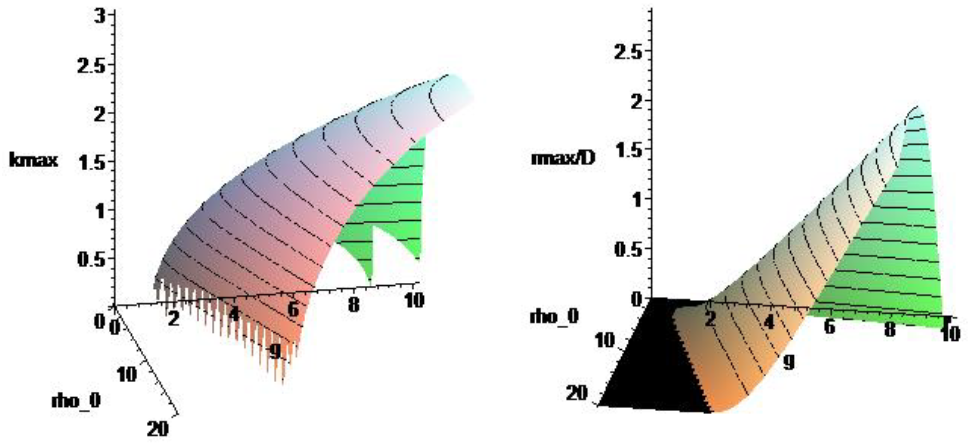
Most rapidly growing wavenumber (*k*_max_) and associated exponential growth rate *r*_max_ (normalized by *D*) as a function of *g* and *ρ*_*0*_ for linear channel gating.

We can see that the dominant feature size of the spontaneously generated pattern should be comparable to the length scale x_0_, although it becomes shorter with stronger instability and stretches out longer at the very margin of instability, before the pattern disappears entirely. In the important limit of small leakage conductance G_0Na_ (i.e. the simultaneous limit of large *g* and *ρ*^0^), *k*_max_ grows with (*ρ*^0^*g*)^1/4^ – which will eventually shrink to the size of a single cell regardless of x_0_.

Similarly, the timescale of growth can be shown to be comparable to τ_P_ ≡ τ_E_ / *k*_*max*_^2^*D* = τ_D_ · (x_0_ / Δx)^4^ / *k*_*max*_^2^, the diffusive timescale τ_D_ scaled by a factor relating the different length scales. Pattern amplitude builds up exponentially on this timescale until it reaches a magnitude comparable to V^*^, at which point it saturates and begins nonlinear evolution. (It is worth noting that the resulting nonlinear evolution and maturation may continue on far longer timescale.) In the limit of small G_0Na_, *k*_*max*_^2^ grows with √(*ρ*^0^*g*), partly offsetting the growth in the (x_0_ / Δx)^4^ factor.

As noted before, chemical kinetics dampen the pattern even as they change its character in interesting ways. τ_K_^−1^ ≡ -*f*’ (*ρ*^0^) / τ_E_ represents the marginal decay rate of ligand concentration due to chemical kinetics, and this is directly subtracted from what would otherwise be the kinetics-free pattern growth rate, as seen above in the equation for *r*. Thus, as a general matter, the ligand decay timescale must be slower than the pattern growth timescale in order for a pattern to form.

### Key criteria for experimental detection

In order for auto-electrophoretic patterning to be detected in a synthetic system or to be offered as an explanation for an existing patterning process, several criteria based on the above analysis must be satisfied:

1. 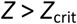 For a pattern to form at all, the combination of ligand and channel must be capable of forming patterns, and the cell must be electrically configured in a favorable manner. Favorable contributing factors include highly charged ligands, cooperative binding, low leakage (ungated) current, large reversal potentials, and ion-selective channels. Background concentration of ligand must be sufficient to have a significant impact on channel current, though not so high as to produce saturation.
2. 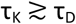 For a pattern to form in the presence of intracellular production and decay of the ligand, the characteristic timescale of the chemical kinetics must be slower than the timescale of pattern formation. This rules out any potential ligands that are well buffered and tightly regulated. If the timescale of pattern formation is hours or days, so must the timescale be for production / decay.
3. 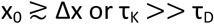 Given the finite size of cells and the high intracellular conductivity compared to gap junction conductivity, the characteristic length scale of pattern formation should be larger than a cell. If it is not, any predicted pattern may be blurred away. In the absence of significant chemical kinetics, numerical experiments show that pattern formation typically still occurs, but on a larger scale.
4. 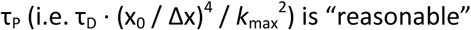 The overall timescale of pattern formation must fit within a reasonable window for experimental observation and must be faster than any un-modeled effects that might disrupt it, such as cell migration. It is easy to pose hypothetical configurations that exhibit interesting pattern dynamics but only on a timescale of days, weeks, or longer. Pattern amplitude is not likely to pose challenges for experimental detection, as the voltage amplitude can be expected to stabilize at a substantial fraction of V^*^, the overall voltage scale of the cell.

### Numerical examples

We now explore some numerical demonstrations of the auto-electrophoresis model, solved using a simple custom code on a hexagonal lattice. Voltage and diffusion are solved implicitly via backward Euler, while advection and chemical kinetics are solved with forward Euler, with ligand conservation enforced via a finite volume style formulation and upwind differencing for advection. Results are not strongly sensitive to time step size, and large time steps are feasible with this scheme, here set to be 0.05τ_P_ in each case.

Above the critical threshold for instability, uniformly distributed ligand will spontaneously form a rippled pattern at the characteristic wavelength. In the absence of chemical kinetics, this will condense into droplet-like aggregates. These aggregates will “coarsen” over long periods of time, smaller aggregates disappearing or merging into larger ones. This is illustrated below for a 100×100 hexagonal mesh:

**Figure 4.**
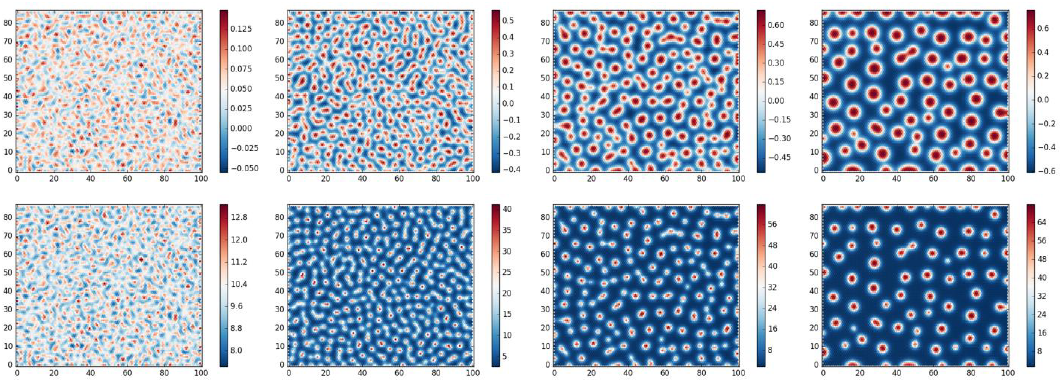
Time points 10τ_P_, 50τ_P_, 250τ_P_, 1250τ_P_. Top frame shows *V*, bottom frame shows *ρ*. *D* = 10^−4^, Δx / x_0_ = 0.25, *g* = 10, *ρ*^0^ = 10, *Z* = 4. τ_P_ ≈ 25τ_D_ = 10^3^τ_E_. Initial conditions are *V* = 0, *ρ* = 10 with uniformly distributed noise at 10% of total amplitude. As anticipated, the pattern amplitude grows rapidly for the first several dozen τ_P_, after which it saturates at a substantial fraction of full scale *V* (−1 to 1) and begins to coarsen on a longer, exponential timescale. This is similar to observations of blowup-free Keller-Segel systems. The particular choice of *D* (and hence the ratio of electric to diffusive timescales) has little impact on the behavior.

In development, these sort of aggregation dynamics could be especially useful for spontaneous polarization. In a small, elongated domain, there will be only two stable steady states: an aggregate adhering to the left wall and an aggregate adhering to the right wall. Multiple aggregates will not be stable, because the larger will eventually subsume the smaller. An aggregate in the middle will also not be stable, because closed boundaries act mathematically like a mirror: the “largest” possible clump is actually a half clump up against the wall, which then appears twice as big. Progressive coarsening thus provides a robust failsafe for spontaneous polarization, able to resolve twinned and malformed states even if the natural wavelength of the initial instability isn’t perfectly matched to the system size.

In the presence of chemical kinetics, one typically finds stable patterns emerging with a definite scale. A wide spectrum of patterns form even with simple linear kinetics – zeroth order production, first order decay. Below shows the formation of a striped pattern (also using a longer length scale):

**Figure 5.**
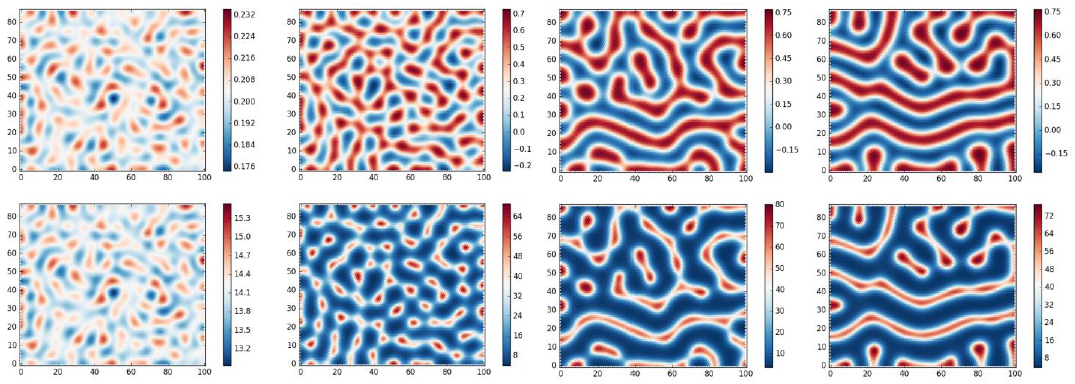
Time points 10τ_P_, 50τ_P_, 250τ_P_, 1250τ_P_. *D* = 10^−5^, Δx / x_0_ = 0.1, *g* = 10, *Z*=4. τ_P_ ≈ 1000τ_D_ = 10^4^τ_E_. k_P_ = 9·10^−5^ ≈ 0.9 / τ_P_, k_D_ = 4·10^−6^ ≈ 0.04 / τ_P_. Initial conditions are *V* = 0, *ρ* = 10 with uniformly distributed noise at 10% of total amplitude.

The variety of patterns appears somewhat richer with longer length scales relative to cell size (x_0_ / Δx), although the time required grows dramatically. With more complex chemical kinetics, patterns of similar characters are seen, but sometimes with nontrivial temporal dynamics, including chaos. *D* again has little impact on the spatial character of the patterns, although larger *D* may sometimes be necessary for complex temporal dynamics.

By varying the parameters of the chemical kinetics across space, one can produce a qualitative map of the differing steady-state pattern characteristics available at different parameters. The variety of patterns nearby the above example are illustrated in the map below.

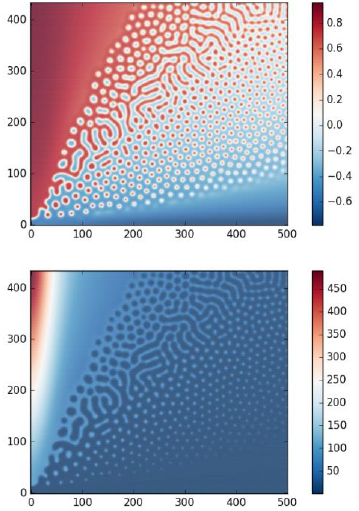

*D* = 10^−5^, Δx / x_0_ = 0.1, *g* = 10, *Z* = 4. 0 ≤ k_P_ ≤ 1.5·10^−4^ on the vertical axis, 0 ≤ k_D_ ≤ 8·10^−6^ on the horizontal axis.

A similar map for the coarser but faster length scale in first example is shown below:

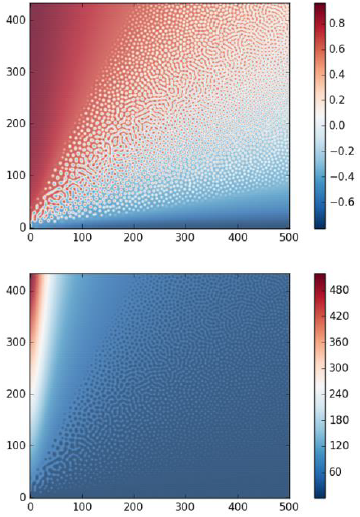

*D* = 10^−4^, Δx / x_0_ = 0.25, *g* = 10, *Z* = 4. 0 ≤ k_P_ ≤ 7.5·10^−4^ on the vertical axis, 0 ≤ k_D_ ≤ 4·10^−5^ on the horizontal axis.

Similar patterns are seen farther to the upper left, gradually weakening in intensity as the chemical kinetics overwhelm the auto-electrophoretic instability.

While these pattern-forming dynamics can potentially be slow, they represent a stable attractor of the system, and thus can be used to refine imperfect hints from prior steps such maternal-effect pre-patterns or patterns generated by high-speed, feed-forward patterning mechanisms. Such a refinement process is dramatically faster than spontaneous generation and can allow the construction of robust, self-healing patterns in a short timeframe.

## References

Brodsky, Micah. 2016. “Partial Redundancy and Morphological Homeostasis: Reliable Development through Overlapping Mechanisms.” Artificial Life 22 (4):518–536. doi: 10.1162/ARTL_a_00216.

Brodsky, Micah Z., and Michael Levin. 2018. “From Physics to Pattern: Uncovering Pattern Formation in Tissue Electrophysiology.” Conference on Artificial Life (ALIFE 2018).

Budrene, E. O., and H. C. Berg. 1995. “Dynamics of formation of symmetrical patterns by chemotactic bacteria.” Nature 376 (6535):49–53. doi: 10.1038/376049a0.

Esser, A. T., K. C. Smith, J. C. Weaver, and M. Levin. 2006. “Mathematical model of morphogen electrophoresis through gap junctions.” Dev Dyn 235 (8):2144–59. doi: 10.1002/dvdy.20870.

Hillen, T., and K. J. Painter. 2008. “A user’s guide to PDE models for chemotaxis.” Journal of Mathematical Biology 58 (1):183. doi: 10.1007/s00285-008-0201-3.

Hiscock, T. W., and S. G. Megason. 2015. “Mathematically guided approaches to distinguish models of periodic patterning.” Development 142 (3):409–19. doi: 10.1242/dev.107441.

Kaupp U. Benjamin, and Reinhard Seifert. 2002. “Cyclic Nucleotide-Gated Ion Channels.” Physiological Reviews 82 (3):769–824. doi: 10.1152/physrev.00008.2002.

Marcon, L., and J. Sharpe. 2012. “Turing patterns in development: what about the horse part?” Current opinion in genetics & development 22 (6):578–84. doi: 10.1016/j.gde.2012.11.013.

Painter, Kevin J., and Thomas Hillen. 2011. “Spatio-temporal chaos in a chemotaxis model.” Physica D: Nonlinear Phenomena 240 (4):363–375. doi: https://doi.org/10.1016/j.physd.2010.09.011.

Pietak, Alexis, and Michael Levin. 2016. “Exploring Instructive Physiological Signaling with the Bioelectric Tissue Simulation Engine.” Frontiers in Bioengineering and Biotechnology 4 (55). doi: 10.3389/fbioe.2016.00055.

Rabinovich, M. I., A. B. Ezersky, and P. D. Weidman. 2000. The Dynamics of Patterns: World Scientific.

Ruiz, M., R. L. Brown, Y. He, T. L. Haley, and J. W. Karpen. 1999. “The single-channel dose-response relation is consistently steep for rod cyclic nucleotide-gated channels: implications for the interpretation of macroscopic dose-response relations.” Biochemistry 38 (33):10642–8. doi: 10.1021/bi990532w.

Turing, A. M. 1952. “The Chemical Basis of Morphogenesis.” Philosophical Transactions of the Royal Society of London. Series B, Biological Sciences 237 (641):37–72. doi: 10.1098/rstb.1952.0012.

Valiunas, V., Y. Y. Polosina, H. Miller, I. A. Potapova, L. Valiuniene, S. Doronin, R. T. Mathias, R. B. Robinson, M. R. Rosen, I. S. Cohen, and P. R. Brink. 2005. “Connexin-specific cell-to-cell transfer of short interfering RNA by gap junctions.” J Physiol 568 (Pt 2):459–68. doi: 10.1113/jphysiol.2005.090985.

Watanabe, M., and S. Kondo. 2015. “Is pigment patterning in fish skin determined by the Turing mechanism?” Trends in genetics: TIG 31 (2):88–96. doi: 10.1016/j.tig.2014.11.005.

Zheng, Q., and G. W. Wei. 2011. “Poisson-Boltzmann-Nernst-Planck model.” J Chem Phys 134 (19):194101. doi: 10.1063/1.3581031.

